# Extracellular Matrix-Guided Tumor Stratification and Network Models Reveal Clinical Molecular Grades

**DOI:** 10.1101/2025.08.29.672994

**Authors:** Aslı Dansık, Sevgi Sarıca, Ece Öztürk, Nurcan Tuncbag

## Abstract

The extracellular matrix (ECM) critically shapes tumor fate and treatment outcome, serving as a potent prognostic factor. Yet, its compositional heterogeneity across tumors makes it difficult to assess its impact on tumor dynamics. To address this, we introduce an ECM-guided patient stratification pipeline through integration of multi-omic data in lung cancer patients. We obtained four patient groups, representing ECM-grades that showed distinct clinical features, mutation profiles, and cellular heterogeneity. Investigation of patient-specific ECM-induced intracellular signaling via network modeling revealed strong enrichment of pathways and transcriptional regulators related to epithelial-mesenchymal transition (EMT) and cancer stemness in higher ECM-grades. Drug proximity analysis on ECM-grade specific networks predicted olaparib as an ECM-grade dependent therapeutic while erlotinib to be ECM-insensitive which were validated experimentally on lung tumor cells with distinct mutational profiles in response to differing ECM microenvironments. Overall, our ECM-mediated stratification approach is a robust system for capturing ECM heterogeneity and identifying patient groups that can be selectively targeted by distinct therapeutic strategies.

## INTRODUCTION

The extracellular matrix (ECM) is an essential regulator of tissue architecture and cellular behavior^1^. It acts as a dynamic reservoir of bioactive molecules, providing both biochemical and mechanical cues that shape cell fate and function^2–4^. Through continuous bidirectional communication with surrounding cells, the ECM undergoes active remodeling, altering its composition and mechanical properties to form tissue-specific microenvironments^5,6^. In tumors, profound alterations of ECM critically shape cancer progression and treatment outcomes^7^ by triggering various processes such as invasion, angiogenesis, epithelial-to-mesenchymal transition (EMT) and metastasis^8–14^. Among them, reduced proteoglycan expression in the ECM leads to better overall survival in patients with lung cancer^15,16^. Elevated expression of cell instructive ECM moieties can induce immune suppression and therapeutic resistance^17,18^. The ECM interacts not only with cancer cells but also with stromal populations such as the cancer-associated fibroblasts (CAFs) and immune cells within the tumor microenvironment (TME) which leads to tumor growth and progression^19–22^. Thus, it is essential to investigate the cellular heterogeneity of the TME and the reciprocal relationship between diverse cellular populations and the altered ECM characteristics^23^. The Matrisome Project disclosed the genes belonging to the “matrisome”, encoding the ECM component proteins and associated factors^24,25^.

Remodeling of the ECM in the TME is highly heterogeneous across patient tumors. Puttock et al. revealed five different ECM composition groups by performing hierarchical clustering on the proteomic data of ovarian cancer samples^26^. Similarly, Bergamaschi et al. found four ECM subgroups in breast carcinomas by performing differential expression analysis on a set of ECM genes, further demonstrating the clinical relevance of these groups^27^. Pankova et al. revealed three subtypes for dedifferentiated liposarcomas (DDLPS) by clustering matrisome/adhesome protein expressions^28^. Parker et al. showed two matreotypes in the squamous cell carcinoma subtype of non-small cell lung cancer (NSCLC) based on expression levels of the ECM genes, with one matreotype having low expression and the other with higher expression and further investigated the differences in these matreotypes^29^. These studies underscored the importance of characterizing ECM heterogeneity and its impact on clinical outcomes.

Although alterations and heterogeneity in tumor matrisomes have been studied extensively, the underlying molecular mechanisms have not been yet described holistically and were limited to single omic layers in previous studies. Network-based methods reveal these mechanisms by identifying latent proteins and pathways by integrating single or multi-omic data layers with their connections^30–32^. These methods are highly utilized in many concepts including novel biomarker identification, drug discovery and repurposing, and patient stratification, yet no such application has been performed with an ECM focus^33–35^. The matrisome landscape is a critical determinant of differential therapeutic response. However, ECM-targeting strategies are mainly limited to targeting of ECM components, cell-ECM interfaces (integrins), or ECM-synthesizing cells (CAFs)^11,36^. Here, network-based methods can help identify potential therapeutic target proteins, revealed as latent factors, beyond the ECM components.

In this study, we developed an ECM-based integrative framework to stratify non-small cell lung adenocarcinoma (LUAD) tumors using multi-omic data. Using the expression profiles of the matrisome components, we defined ECM barcodes for each patient and classified tumors into distinct ECM grades (ECM-G1 … ECM-Gn), reflecting their differential enrichment. In parallel, we constructed patient-specific networks and derived consensus networks characterizing each ECM grade. Computational drug screening based on network proximity repurposed several candidate drugs with potential ECM-grade-specific activity, linking molecular networks to ECM-stratified therapeutic options. Finally, we experimentally validated the identified hits on LUAD cell lines with distinct mutation profiles grown in interaction with differing ECM that represent healthy and tumor microenvironments, revealing ECM-dependent and independent therapeutics (Figure 1).

**Figure 1.**
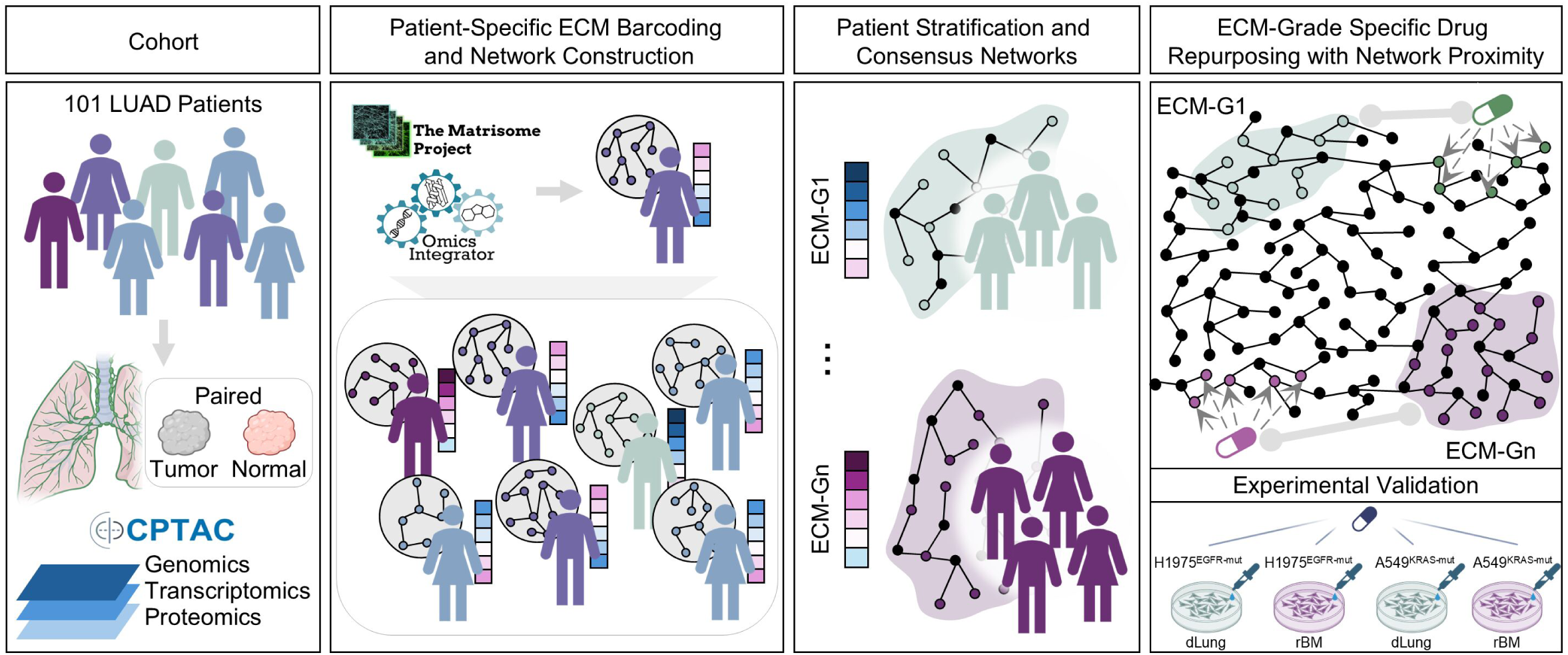
Overview of patient stratification, network modeling and drug repurposing. Multi-omic data of tumors and normal samples of 101 patients with LUAD were retrieved from CPTAC. Each patient was barcoded based on the ECM gene expression belonging to categories retrieved from The Matrisome Project. Next, patient-specific networks were constructed with Omics Integrator by integrating phosphorylated ECM proteins and transcriptional factors regulating highly dysregulated ECM genes in tumors compared to matched normal tissue. Then, an ECM grade was assigned to each patient, and a consensus network was constructed for each ECM grade, which was used for network-proximity-based analysis to rank drugs that are potentially ECM-grade specific. Finally, experimental validation of ECM-dependent drug response was performed using LUAD cell lines cultured in healthy and tumorigenic microenvironments.

## RESULTS

### ECM-based patient stratification reveals both shared and distinct matrix signatures

Aberrant ECM remodeling is a hallmark of tumor progression and may distinguish malignant from normal tissue^37^. To explore this in the CPTAC LUAD patient cohort (n=101), we first conducted unsupervised transcriptomic and proteomic analyses of the matrisome from the tumors and matching normal tissues. Out of 1028 matrisome genes 951 were detected in the transcriptomic data leading to a coverage of 92%. The distribution of expressed ECM genes across divisions (core and associated) and categories (collagens, glycoproteins, proteoglycans, ECM regulators, ECM-affiliated proteins and secreted factors) closely matched those defined by the Matrisome Project^25^, indicating a representative coverage of ECM components in this patient cohort (Figure 2a). The categorical distribution of ECM genes in proteomic data was also consistent with the transcriptomic data with marginal deviations (Supp Fig 1a). In addition, we found that the expression of ECM genes and their protein products were significantly correlated both globally (ρ_spearman_=0.60, p-value<0.001) and within individual patients (median [Q1–Q3] ρ_spearman_ : 0.56 [0.50–0.61]) (Supp Fig 1b, c). Hierarchical clustering of ECM gene expression profiles of all samples revealed a clear separation between tumors and matching normal samples (Supp Fig 1d). Principal component analysis (PCA) showed ECM-dependent distinction between tumor and normal samples with two well-separated clusters. Notably, we observed increased transcriptional heterogeneity of ECM genes in tumors displaying higher intra-group variance (avgED_tumor_=15.57) compared to normal samples (avgED_normal_=8.038) (Supp Fig 1e). This pattern and intra-group variances of ECM protein levels were preserved when the proteomic data was used for the comparison (avgED_tumor_=23.98, avgED_normal_=13.02) (Supp Fig 1f, g).

**Figure 2.**
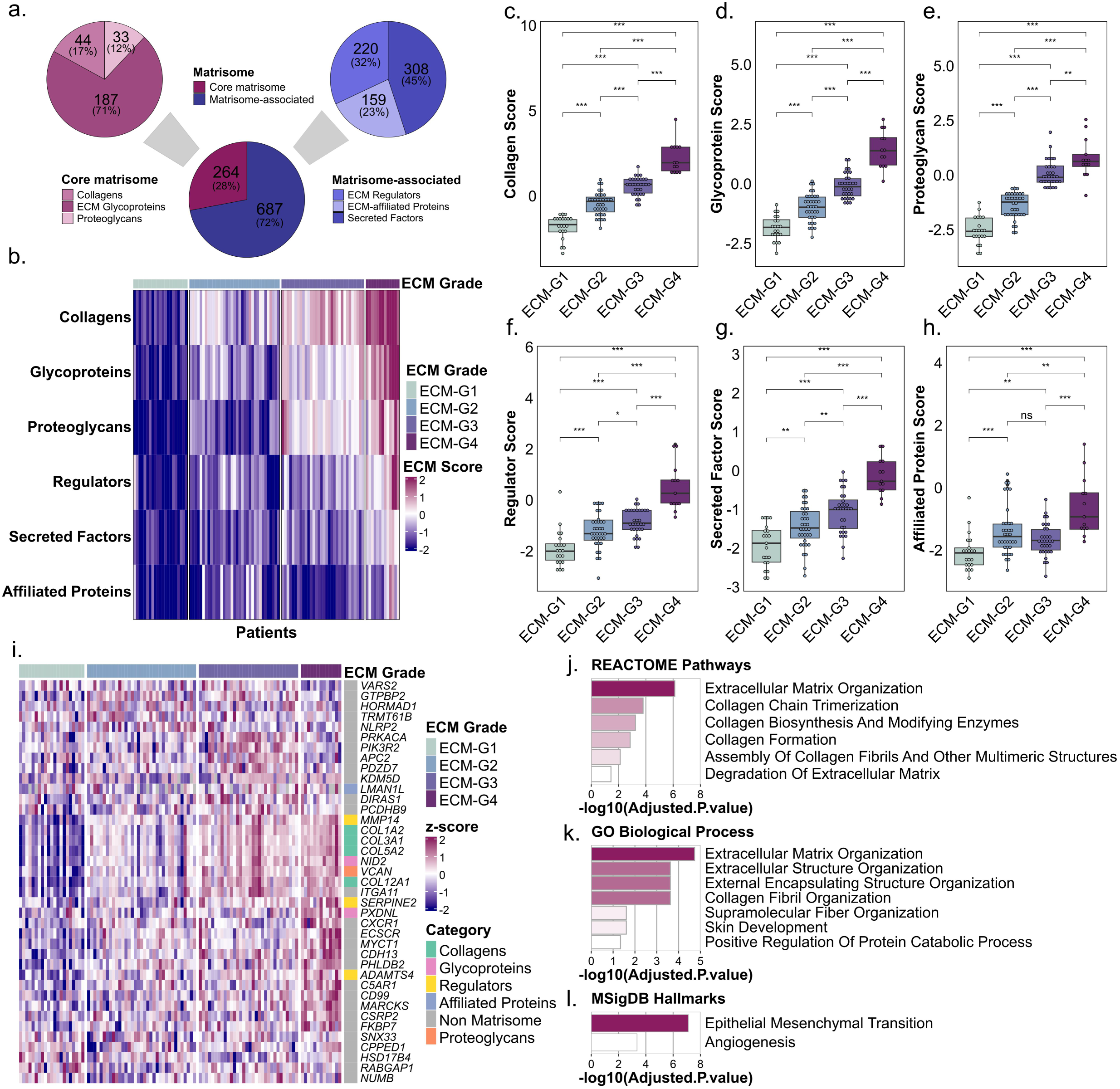
Multi-omic ECM barcodes reveal four consensus clusters that both dependently and independently represent the ECM grades of patients. **a**. Proportions of the expressed matrisome divisions (core, matrisome-associated) and categories (left panel: collagens, glycoproteins, proteoglycans; right panel: ECM regulators, ECM-affiliated proteins, and secreted factors) in the CPTAC cohort. **b.** The heatmap shows the four identified ECM grades in ascending order from left to right where each grade is indicated with a unique color shown at the top. Each column represents a patient, and each row represents an ECM category score that indicates the multi-omic expression of ECM genes. **c–h.** Boxplots show differences in ECM scores across ECM grades. Each box extends from the lower to the upper quartile where the horizontal line indicates the median value. Statistical analysis was Mann– Whitney U test, two-sided, ns: non-significant, p*<0.05, p**<0.01, p***<0.001. **i.** Expressions of highly variable genes independently identified across ECM grades are shown in the form of z-scores. The sidebar shows the ECM categories of genes with distinct colors where non-matrisome genes are indicated with grey. **j–l.** Enriched REACTOME pathways, GO biological processes, and MSigDB Hallmark pathways identified from functional enrichment analysis performed on the highly variable genes (Adjusted.P.Value < 0.05).

To investigate this variation, we stratified the patients based on their ECM gene expression profiles. Here, we computed scores for six ECM categories for each patient by integrating transcriptomic and proteomic expression data which reflects the average expression of genes within each category. The resulting six-category molecular signature was referred to as the patient’s ‘ECM barcode’, providing a compact representation of its category-specific expression profile. Next, we applied reference-based consensus clustering and found the number of clusters as four (Monte Carlo p-value<0.05) which was the optimal solution based on parameter tuning (Supp Fig 1h-j). The resulting distinctive gradient-like pattern across the four patient clusters clearly reflected a progression from low grade to high grade ECM enrichment (Figure 2b). Thus, we labeled the patient groups as ECM Grades 1 to 4 (ECM-G1, ECM-G2, ECM-G3, ECM-G4). PCA positioned ECM-G1 and ECM-G4 at opposite ends of the first principal component (PC1), with ECM-G2 and ECM-G3 distributed between them, indicative of transitional states (Supp Fig 1k).

ECM-G1 (n=21) with the lowest grade of ECM richness presented almost no enrichment in ECM categories but only a very mild expression in collagens compared to other groups. ECM-G2 (n=35) was the second-least ECM-enriched group but was more enriched in collagen and glycoproteins when compared with ECM-G1. ECM-G3 (n=32) was again more enriched in collagens and glycoproteins, but additionally in proteoglycans, when compared with ECM-G2. Lastly, ECM-G4 (n=13) showed the highest enrichment in all ECM categories. Collagen content was the common ECM category that was enriched in the majority of ECM grades at different levels, which was not surprising since collagen is highly expressed in many types of solid tumors^38^. We observed a statistically significant increase in all matrisome component scores with increasing ECM grades. Notably, ECM-affiliated proteins exhibited no significant change between intermediate grades (Figure 2c–h).

To independently validate that the ECM grades reveal alterations in ECM characteristics, we identified highly variable genes between these groups across the entire transcriptomic data and found a set of 39 genes, which were significantly enriched in matrisome genes (Overlap = 11 genes, Fisher’s Exact Test p-value<0.001) (Figure 2i). As a result of the functional enrichment analysis, we found that the top ranking REACTOME pathway was ECM organization, followed by pathways related to the synthesis, modification, formation, and degradation of ECM or its components (Figure 2j). Similarly, the most enriched GO biological process was ECM organization, followed by processes related to external structure and fibril organization (Figure 2k).

Additionally, genes and hallmark pathways related to metastasis, angiogenesis, and apoptosis were significantly enriched. ITGA11, a collagen receptor whose overexpression in NSCLC has been associated with tumor growth, metastasis, and reduced recurrence-free survival, also showed increased expression with elevated ECM grade^39^. PIK3R2, a downstream mediator of FAK that can lead to AKT activation and subsequent inhibition of apoptosis, was highly expressed in ECM-G3^40^. Hallmark pathways of EMT and angiogenesis were also highly enriched, demonstrating the significant influence of ECM profiles on such processes (Figure 2l). Collectively, our ECM-dependent and - independent analyses showed that the identified ECM grades effectively captured key population-level ECM features, delineated distinct expression profiles, and were significantly associated with metastasis and apoptosis inhibition.

### Higher ECM grades are characterized by enhanced cellular heterogeneity, increased EMT activation and invasive potential

To validate that our ECM-based stratification framework captures the cellular composition and heterogeneity associated with each grade in the TME, we performed cellular deconvolution on the transcriptomic data. We found that distinct grades were characterized by specific patterns of cell type enrichment and diversity (Figure 3a). In ECM-G4, tumors exhibited, on average, a 4.89-fold increase in CAF content compared to their normal counterparts. However, CAF content was highly variable in ECM-G1 tumors (median [Q1– Q3] log2FC : - 0.064 [-0.96–0.32]) and did not reflect a distinctive pattern. We observed a significant increase in CAF enrichment with elevation in ECM grade, suggesting that CAF content is highly sensitive to ECM profiles (Figure 3b). Endothelial cell enrichment was reduced in tumors across all ECM grades compared to their normal counterparts; however, the loss was more pronounced in ECM-G1 and significantly less in ECM-G4, suggesting that the TMEs in ECM-G4 may be more conducive to maintaining endothelial cell growth (Figure 3c). Higher ECM deposition likely present in this group may lead to hypoxic conditions, which enable the recruitment of endothelial cells. To cross-check our findings, we performed cellular deconvolution using another tool called EPIC and observed similar results (Supp Fig. 2a, b). Unlike CAFs and endothelial cells, we observed no ECM-dependent difference in the enrichment of immune-related cell populations.

**Figure 3.**
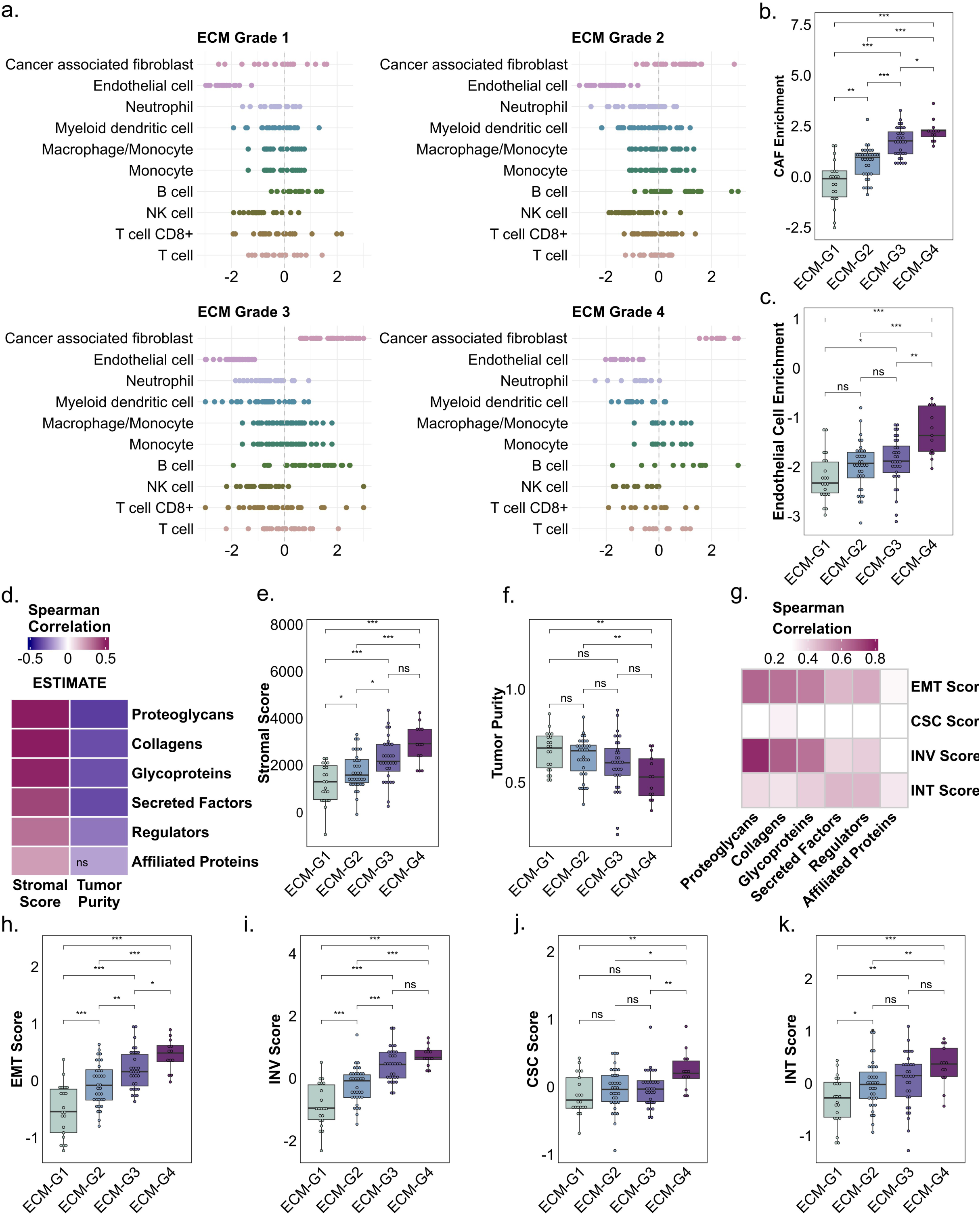
ECM grades show altered stromal cell maintenance promoting EMT and invasiveness features. **a.** Cellular deconvolution of gene expression data reveals enrichment scores for 10 major cell populations, including CAFs, endothelial cells, and immune cells. For each cell type, the log2 fold change in enrichment in patient tumors compared to matched normal tissues is calculated for each ECM grade. **b, c.** Boxplots show differences in CAF and endothelial cell enrichment across ECM grades. **d.** The heatmap shows the correlation between ECM scores in the rows and stromal scores and tumor purity in the columns (Spearman’s rank correlation, ns: non-significant). **e, f.** Boxplots show the difference in stromal score and tumor purity across ECM grades. **g.** The heatmap shows the correlation between ECM scores in the columns and EMT, CSC, INV, and INT scores in the rows (Spearman’s rank correlation, ns: non-significant). **h-k.** Boxplots show the difference in EMT, CSC, INV, and INT scores across ECM grades. All statistical tests used in boxplots are Mann–Whitney U test, two-sided, ns: non-significant, p*<0.05, p**<0.01, p***<0.001. Each box extends from the lower to the upper quartile where the horizontal line indicates the median value.

Next, we examined overall stromal cell content enrichment and the proportion of tumor cells using the ESTIMATE stromal score and tumor purity, respectively. ESTIMATE stromal scores showed positive correlations with the ECM scores, while tumor purity was negatively correlated as expected (Figure 3d). All the ECM scores showed a significantly positive correlation with the stromal score, with the lowest being the affiliated proteins (ρ_spearman_=0.22, p-value<0.05). Especially core matrisome genes were highly correlated, with the highest being the proteoglycans (ρ_spearman_=0.58, p-value<0.001). Tumor purity, on the other hand, showed a significantly negative correlation with the ECM scores, except for the affiliated proteins, with the strongest negative correlation observed for the proteoglycans (ρ_spearman_=-0.37, p-value<0.001). We also compared the changes in stromal score and tumor purity between ECM grades and observed a significant increase in stromal score with increasing ECM grades, except between ECM-G3 and ECM-G4 (Figure 3e). The tumor purity of ECM-G4 was significantly lower than that of ECM-G2 and ECM-G1, with an average purity of 0.53, while ECM-G1 showed the highest average purity at 0.67 (Figure 3f).

Collectively, these results show that the abundance of stromal cells in the TME increased as the complexity and abundance of the ECM increased—resulting in a more developed TME. Lower tumor purity we observed in higher ECM grades can be associated with poor prognosis since a higher population of stromal cells is known to support tumor growth and development^22,41,42^. These stromal cells, especially CAFs, also guide other cell types, such as endothelial cells, to proliferate, which explains the increase we observed for the endothelial cells along with CAFs^23^. This positive reinforcement between the ECM grades and stromal cell content gives rise to more developed and higher-risk tumors.

Next, we calculated average expression scores of various cancer-associated gene groups like EMT, cancer stem-cell (CSC) related, invasiveness (INV), and integrins (INT), to investigate the influence of ECM enrichment. We observed a significant correlation of core matrisome scores with both EMT and INV scores (Figure 3g). The proteoglycan score was the most positively correlated with both EMT (ρ_spearman_=0.64, p-value<0.001) and INV (ρ_spearman_=0.77, p-value<0.001) scores, consistent with our previous findings solely obtained from transcriptomic data^16^. We also observed a significant increase in EMT and INV scores with increasing ECM grades (Figure 3h, i). Unlike these scores, we observed a low correlation between ECM scores and CSC and INT scores. Only the collagen score was moderately correlated with CSC (ρ_spearman_=0.33, p-value<0.001) score, in line with the maintenance of CSC self-renewal by collagen^43^. ECM-G4 had a significantly higher CSC score when compared to the other grades, suggesting that the maintenance of CSCs could require very aggressive ECM profiles (Figure 3j). ECM-G1 showed significantly reduced INT scores when compared with other groups except for ECM-G2, yet the remaining groups showed similar INT scores (Figure 3k). Our results suggest that a moderate or high ECM enrichment is necessary for ECM-integrin interaction, and when ECM profiles are poor, such interactions could not be facilitated as efficiently.

### ECM grades associate with divergent clinical features and exclusive oncogenic mutations

We next investigated the survival and recurrence probabilities of patients with distinct ECM grades. Survival analysis showed that patients stratified in ECM-G4 group had poor overall survival, and ECM-G1 had the best survival probability among them, where ECM-G2 and ECM-G3 showed moderate survival. We also observed a similar trend for recurrence-free patterns, where ECM-G4 showed a more rapid decline in recurrence-free proportion compared to other groups, and ECM-G1 presented a lower recurrence probability (Figure 4a). To investigate the rapid decline at earlier time points in both curves, we performed a short-term analysis, revealing a significant decrease in survival and recurrence-free probabilities between ECM-G1 and ECM-G4 within the first 600 and 300-day periods, respectively (the log-rank test, Figure 4b). These findings indicate that patients with more enriched ECM exhibit poorer survival and a higher probability of recurrence than patients with less enriched ECM. The high recurrence rate in the ECM-rich group may stem from the exclusively higher enrichment of CSC markers, well-known drivers of recurrence^44^. To further evaluate the prognostic relevance of ECM grades, we performed Cox proportional hazards regression analysis across the patient cohort. Patients with ECM-G4 exhibited a six-fold higher risk of death compared to the patients with ECM-G1, suggesting a strong association between higher ECM grades and poor survival outcomes (Figure 4c).

**Figure 4.**
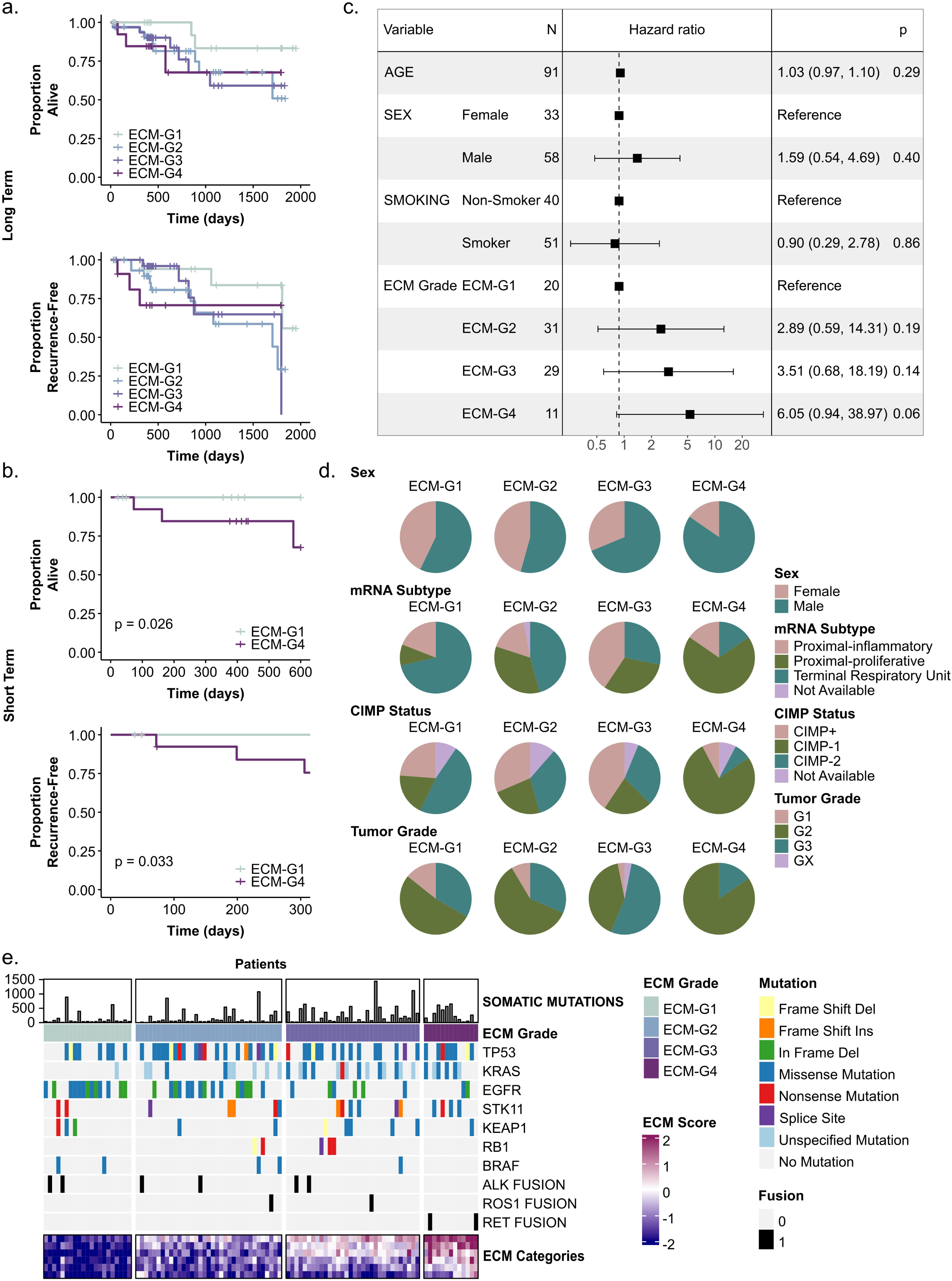
ECM grades reveal grade-dependent prognosis, clinical features and mutual exclusivity. **a, b.** Kaplan-Meier plots of ECM grades showing proportion alive and recurrence-free in long- and short-term periods (log-rank test). **c.** Multivariate hazard ratio plot taking into account age, sex, smoking status, and ECM grade (Cox proportional hazards regression). **d.** Distribution of patients in ECM grades according to their sex, mRNA subtype (PI, proximal inflammatory; PP, proximal proliferative; and TRU, terminal respiratory unit), CIMP status (CIMP+: highest methylation, CIMP-1: high methylation, CIMP-2: moderate methylation), and tumor grade. **e.** The heatmap displays the mutation landscape of cancer driver genes and mutation burden across ECM grades. The top panel indicates the number of somatic mutations per patient, while the lower panels highlight mutations and gene fusions in established driver genes, with mutation types color-coded and fusions marked in black. The bottom panel presents ECM scores.

To explore potential clinical patterns across the ECM groups, we examined additional clinical variables and observed trends in the distribution of factors such as sex, mRNA subtypes, CIMP status, and tumor grade (Figure 4d). mRNA subtypes represent distinct expression programs: PI (proximal inflammatory) is characterized by immune-related and inflammatory gene expression; PP (proximal proliferative) by elevated cell cycle and proliferation signals; and TRU (terminal respiratory unit) by gene expression of functional respiratory epithelium. CIMP status provides information on the extent of methylation within the CpG islands (CIMP+: highest methylation, CIMP-1: high methylation, CIMP-2: moderate methylation). ECM-G4 had the highest percentage of males (84.61%) among the grades, and it has been shown that males have higher risk and mortality compared to females in lung cancer^45^. Consistently, ECM-G4 exhibited one of the lowest long- and short-term survival rates, and the hazard ratio for men was higher indicating a 59% increased risk compared to women. Regarding the mRNA subtype, ECM-G4 had the highest percentage of patients having the PP subtype (69.23%) and the lowest percentage of TRU subtype (15.38%), while ECM-G1 had the contrary (9.52% PP and 71.42% TRU). An apparent increase in the proportion of patients with the PP subtype was observed with increasing ECM grade. These results imply a shift in transcriptomic profile from a normal lung tissue to a proliferation-related profile. The PI proportions remained rather constant among all grades, except ECM-G3 (40.62%), suggesting that this group may have an altered immune response. For the CIMP status, ECM-G1 had the highest percentage for the CIMP-2 status (47.61%), while this percentage showed a decrease as the ECM grades increased, and the lowest percentage of CIMP-2 status was observed for ECM-G4 (7.69%). Although a higher percentage of CIMP+ patients was observed in ECM-G1, the overall percentage of CIMP-1 and CIMP+ was much higher in ECM-G4 and ECM-G3. Our results indicate that overall hypermethylation of CpG islands was much more common in ECM-G3 and ECM-G4, making these groups more susceptible to the silencing of tumor suppressors. Finally, tumor grades showed an altered pattern as the ECM grades shifted. ECM-G1 was the group that showed the highest percentage for tumor grade 1 followed by ECM-G2, ECM-G3 and ECM-G4 with percentages 14%, 8.57% 3.12% and 0% respectively. ECM-G3 held the highest percentage for tumor grade 3 (53.12%). Overall, we conclude that tumor grades are also influenced by the ECM grades, and the mildest tumor grade is depleted in ECM-G4.

We then investigated whether mutation profiles and burden of patients changed with the ECM grade (Figure 4e). We observed that the highest mutation load was in ECM-G4 followed by ECM-G3, ECM-G2 and ECM-G1 with median somatic mutation count of 216, 188.5, 55.5, 42, respectively. This provided evidence that somatic mutations accumulate more in ECM-rich groups than in poor ones. High mutation counts are known to give rise to more aggressive cancer phenotypes and higher tumor heterogeneity, which was shown to be the case for ECM grades as they went higher^46,47^. Positive correlation of ECM grades with mutation load suggests a mechanism that leads to genomic instability. Indeed, ECM can directly affect DNA repair mechanisms and influence the probability of somatic mutations by, for instance, ECM-induced stress or by stimulating mechanical signaling pathways by exerting mechanical forces on cells^48,49^.

We also checked the mutation status of tumor drivers among ECM grades. Not surprisingly, TP53 mutation was the most frequent across all ECM grades since it is one of the most mutated genes in LUAD^50^. Yet ECM-G4 (61%) and ECM-G3 (61%) had the largest proportion of patients with TP53 mutation followed by ECM-G2 (46%) and ECM-G1 (38%).

Strong driver mutations of LUAD tumors, KRAS and EGFR, show mutual exclusivity^51^. We observed a higher KRAS mutation accumulating on richer ECM grades (p-value<0.01) and a higher EGFR mutation on poorer ECM grades (p-value<0.001), presenting not only the mutually exclusive nature of these drivers but also how ECM grade influences this exclusivity. Our analysis revealed that poor ECM grades like ECM-G1 and ECM-G2 were more prone to having EGFR mutations while rich ECM grades, including ECM-G3 and ECM-G4, tend to have KRAS mutations, suggesting that the increase in abundance of ECM can compensate for EGFR mutations and stimulate growth-inducing signaling pathways.

### Patient-Specific Networks Reveal ECM Grade–Dependent Activation of EMT, Angiogenesis, and Key Transcriptional Regulators

To investigate the connectivity between ECM grades and causal signaling pathways and to reveal latent molecular patterns and transcriptional regulators, we reconstructed patient-specific networks based on their ECM characteristics. These patient-specific networks preserved inter-tumor heterogeneity while incorporating high-confidence protein interactions (Supp Fig 3a). When patient-specific networks were merged as consensus networks that represent each ECM grade, ECM-G2 exhibited the highest protein coverage (|v|=218), followed by ECM-G1 (|v|=204), ECM-G3 (|v|=203), and ECM-G4 (|v|=172) (Figure 5a). In total, 104 proteins were shared among all groups. While many proteins were shared in at least three ECM grades, only two proteins were uniquely shared between ECM-G1 and ECM-G4. In contrast, consecutive grade pairs (ECM-G(i)-ECM-G(i+1)) exhibited a distinct subset of proteins (47 proteins) shared exclusively between them and Jaccard similarity between consensus networks showed the greatest similarity between ECM-G2 and ECM-G3 (Supp Fig 3b), suggesting molecular convergence between the intermediate tumor states and highlighting the divergence of the lowest and highest ECM grades. This was consistent with the unsupervised separation of ECM grades via PCA (Supp Fig 1k). Despite common proteins and pathways, their expression or activity levels may vary significantly across ECM grades, reflecting ECM-specific regulatory effects rather than sole presence. To capture this effect, we calculated patient-specific network scores by simply averaging the expression of the network proteins and observed a significant ECM-grade dependent elevation of the network scores (Figure 5b). We need to note that these networks contain both matrisome and intermediate proteins involved in intracellular signaling. The monotonic increase in network scores along ECM grades suggests their collective upregulation in higher ECM grades and downregulation in lower ones.

**Figure 5.**
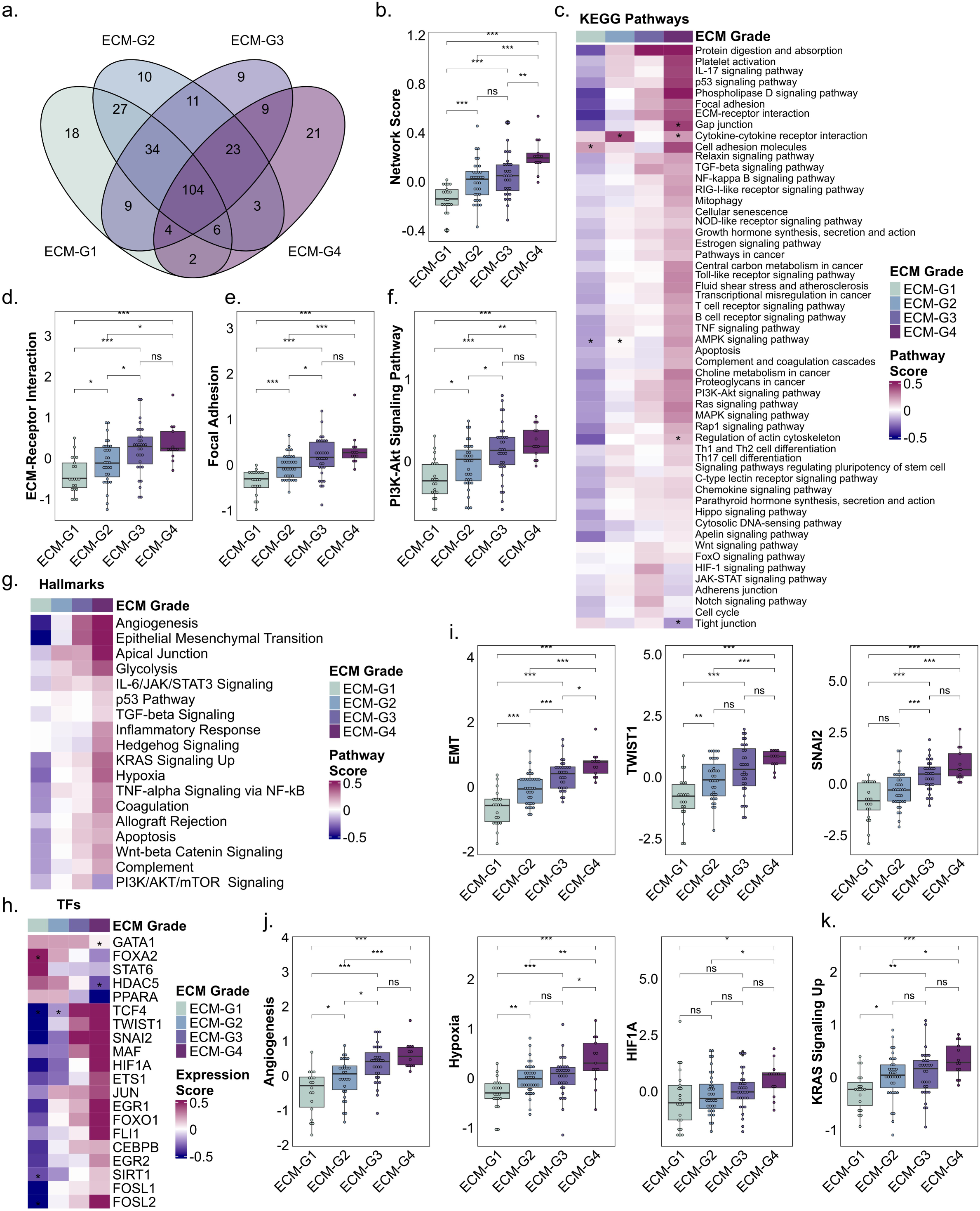
ECM grade-specific enrichment patterns reveal a high impact of ECM on pathways and transcriptional regulators related to tumor growth and metastasis. **a.** Shared and unique proteins of ECM grade-specific consensus networks we derived by merging patient-specific networks are shown. **b.** Boxplot shows differences in network scores across ECM grades which indicate the expression levels of the proteins in patient-specific networks. **c.** The heatmap shows significantly enriched KEGG pathways in patient-specific networks of each ECM grade. “*” shows pathways enriched in less than 25% of the patients in an ECM grade. **d-f.** Boxplots show differences in ECM-receptor interaction, focal adhesion, and PI3K-Akt signaling pathway scores across ECM grades, indicating the average expression of overlapping proteins between patient-specific networks and the given pathway. **g, h.** The heatmap shows significantly enriched MsigDB Hallmark pathways and TFs in patient-specific networks of each ECM grade. “*” shows enrichment in less than 25% of the patients in an ECM grade. Expression scores of the TFs indicate the median expression of the TF. **i.** Boxplots show differences in EMT pathway, TWIST1, and SNAI2 expression scores across ECM grades. **j.** Boxplots show differences in angiogenesis, hypoxia pathway, and HIF1A expression scores across ECM grades. **k.** Boxplots show differences in KRAS signaling up pathway scores across ECM grades. All statistical tests used in boxplots are Mann–Whitney U test, two-sided, ns: non-significant, p*<0.05, p**<0.01, p***<0.001. Each box extends from the lower to the upper quartile where the horizontal line indicates the median value.

We also revealed that several pathways exhibited ECM-grade-dependent enrichment in patient-specific networks such as pathways related to ECM, cell adhesion, proliferation, EMT, metastasis, apoptosis and cell senescence (Figure 5c). Deconvolution of the reconstructed networks into pathway modules revealed that the grade-dependent trend of expression increase was preserved at the pathway level besides the network level.

Many ECM-related pathways, like ECM-receptor interaction, focal adhesion, regulation of the actin cytoskeleton, and proteoglycans in cancer, were enriched for higher ECM grades. ECM-receptor interaction and focal adhesion showed significantly increased enrichment across ECM grades (Figure 5d, e). Focal adhesion complexes mediate cell-ECM interactions, and their dysregulation has been linked to enhanced tumor invasiveness^52–54^. Accordingly, our pathway enrichment analysis suggests increased invasive potential with higher ECM grade and aligns with the significant correlation we observed with the INV score (Figure 3i).

We found a similar trend in the PI3K-Akt signaling pathway (Figure 5f) —a key pathway in mechanosensing— activity^55^. Elevated ECM deposition, likely observed in higher ECM grades, may be exerting mechanical stress on tumor cells, activating this pathway and thereby promoting cell migration and tumor growth. We additionally observed several cancer hallmark pathways such as, p53, TGF-β, TNFα, NFκB, Wnt, and PI3K signaling (Figure 5g) and significantly active key transcriptional regulators (Figure 5h) that show monotonically increasing or decreasing pathway and TF expression scores across different ECM grades.

The EMT hallmark score increased significantly and progressively with ECM grade, accompanied by elevated expression of the canonical EMT transcription factors TWIST1 and SNAI2 (Figure 5i). ETS1, previously associated with poor prognosis and EMT progression^56,57^, also showed increased expression in higher ECM grades (Figure 5h). In contrast, FOXA2, a factor known to correlate with suppression of invasion and TGF-β-induced EMT^58,59^, was markedly downregulated in ECM-rich grades and significantly enriched in ECM-G1 (Supp Fig 3c). In addition, while cell adhesion pathways were increasingly enriched in higher ECM grades, pathways of cell-cell junctions—such as gap, adherens, and tight junctions—did not follow the same pattern (Figure 5c). Enrichment of the tight junction pathway, associated with the epithelial phenotype and EMT suppression^60^, was low in ECM-G4 (below 25%). In contrast, the TGF-β signaling pathway, a key pro-EMT regulator^61^, was increasingly enriched in higher ECM grades. Collectively, these results reveal a positive association between ECM grades and pro-EMT regulators, and a negative association with anti-EMT regulators, consistent with our EMT score-based findings (Figure 3h).

Hallmarks of angiogenesis and hypoxia as well as HIF1A, a regulator of VEGF and a key factor in inducing angiogenesis, were increasingly enriched across ECM grades (Figure 5j)^62^. Increased angiogenesis is a driver of metastatic potential, suggesting that patients with higher ECM grades may be more susceptible to metastasis^63^ – potentially explaining the elevated recurrence rate in these groups (Figure 4a, b).

KRAS signaling was increasingly enriched in higher ECM grades (Figure 5k), in line with the significant accumulation of KRAS mutations in ECM-rich grades (Figure 4e). KRAS-mutant cancer cells can remodel ECM by upregulating metalloproteinases (MMPs)^64^, potentially explaining the positive correlation between KRAS signaling and increasing ECM grades in our analysis. Previous studies have shown that KRAS-mutant lung cancer cell lines with downregulated FOXA2 expression exhibited elevated EMT markers, increased migration, and enhanced metastatic potential, whereas FOXA2 expression in EGFR-mutant lung tumors was associated with suppressed tumor progression^59,65^. This differential role of FOXA2 aligns with KRAS and EGFR mutation profiles and EMT and INV score outcomes observed across ECM grades in our analysis, highlighting a potential mechanistic link between mutational context and ECM grade-dependent invasiveness.

### Drug screening via network proximity reveals ECM-sensitive and insensitive therapeutics

Aberrant changes in the ECM are known to influence response to therapy^17^. To investigate the effects of ECM grades on the efficacy of therapeutics, we performed network-based proximity analysis for drug screening. We used four grade-specific consensus networks. The ECM-G4 network was enriched in various hallmark pathways, such as EMT (Overlap = 19 genes, p-value<0.001), angiogenesis (Overlap = 4 genes, p-value<0.001), hypoxia (Overlap = 14 genes, p-value<0.001), KRAS (Overlap = 10 genes, p-value<0.001) and PI3K signaling (Overlap = 6 genes, p-value<0.001) pathways (Figure 6a). These pathways cover both matrisome proteins and intermediate proteins, suggesting that the modeling revealed latent players that could not be identified by high-throughput technologies. We also observed that the consensus networks preserved the shared pathways present in patient-specific networks of the same ECM grade.

**Figure 6.**
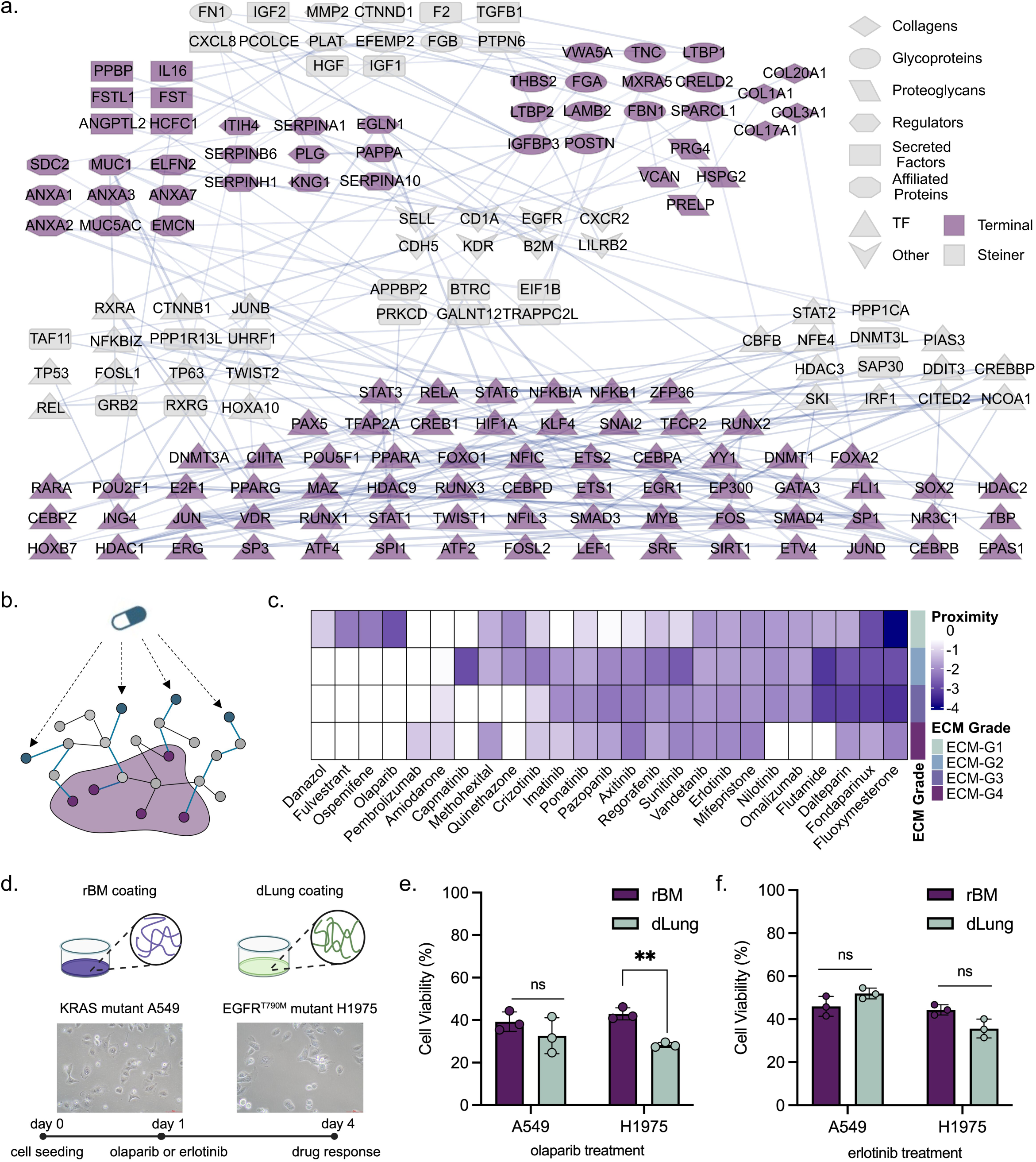
Network-based screening of drugs reveals ECM grade-dependent and independent hits. **a.** Consensus network of ECM grade 4. **b.** Schematic representation of network-based drug screening where blue nodes represent drug targets and purple nodes represent network proteins. **c.** The heatmap shows proximity scores of drugs on consensus networks of ECM grades. **d.** Schematic representation of experimental validation of drugs. **e, f.** The bar plots show the cell viabilities of A549 and H1975 cells in rBM and dlung environments when subjected to olaparib and erlotinib. Three independent replicates were measured, and statistical significance was tested with unpaired t-test, ns: non-significant, p**<0.01.

To identify potential drugs targeting different ECM grades, we conducted network proximity analysis between drug target proteins and the consensus networks of each grade. For each network protein, the shortest path to any drug target was determined, the minimum distance was selected, and the average of these minimal distances was calculated. This average was then compared to a distribution of distances from randomized networks to derive a relative proximity score (𝑧_𝑐_) (Figure 6b). We retrieved target proteins of drugs from DrugBank (in total, 5961 drugs), filtered out those that have a number of targets higher than 25 (after filtering, 5893 drugs), and calculated proximities. Then, only FDA approved high-affinity inhibitor drugs curated from OncoTarget analysis (194 drugs)^66^ were used for further analysis. The candidate drugs were shortlisted based on network-proximity measures where drugs with 𝑧_𝑐_< -2.0 present in at most two groups or having a continuous trend of proximity values across ECM grades (25 drugs). As a result, we revealed both cluster-specific and large-spectrum drugs with possibly higher efficiencies on specific ECM groups (Figure 6c). Pembrolizumab, an approved biotech drug that blocks PD-1 and is used for the treatment of various cancers like metastatic melanoma and NSCLC, was the only drug with a high relative proximity ( 𝑧_𝑐_ =-1.21) to ECM-G4. Capmatinib, an approved inhibitor of c-Met receptor tyrosine kinase (MET) known to be used in NSCLC, was significantly proximate to ECM-G2 (𝑧_𝑐_=-2.92), while it showed no proximity to others. This kinase was only present in ECM-G2, which explains the proximity of this drug to this group. Danazol, fluvestrant, ospemifene, and olaparib were in proximity with ECM-G1 while showing no proximity to others. Olaparib, an approved inhibitor of poly (ADP-ribose) polymerase (PARP) enzymatic activity, known to be used to treat the number of cancers by mainly acting on PARP1 and PARP2, was significantly proximate to this group (𝑧_𝑐_=-2.85)^67^. Both proteins were present in the consensus network of ECM-G1, while only PARP1 was present in ECM-G2, and none of them were present in the remaining groups. This could explain the significantly higher proximity of this drug to ECM-G1. In addition to cluster-specific drugs, we also observed ECM grade-independent drugs that show high proximity to all of the groups, like erlotinib and vandetanib, which are both tyrosine kinase inhibitors mainly targeting EGFR and VEGFR, respectively^68,69^. All in all, this analysis shows that ECM grades may be targeted by specific drug molecules, while some are ECM grade-independent.

We next aimed to experimentally validate the findings of the in-silico network-based drug proximity analysis. We seeded KRAS or EGFR mutant NSCLC cell lines onto different matrices aiming to assess the ECM dependency of drug response. To represent a low-grade ECM microenvironment, we used healthy lung-derived decellularized ECM (dLung)^70^, whereas tumor-derived reconstituted basement membrane (rBM) was used to represent a higher-grade ECM enriched in cell instructive matrix cues^71^. Among the identified drug group, PARP inhibitor olaparib (ECM grade-dependent) and EGFR inhibitor erlotinib (ECM grade-independent) were chosen to test whether given ECM cues regulate the efficacy of these drugs. Illustration of the experimental setup shows the bright-field images of KRAS-mutant A549 cells and EGFR-mutant H1975 cells and the timeline of drug treatment process (Figure 6d). Treatment with olaparib showed an ECM-dependent regulation of response for EGFR-mutant H1975 cells and potency of the drug is higher when cells were seeded onto dLung-coated surfaces (Figure 6e). Intriguingly, KRAS-mutant A549 cells showed similar sensitivity to olaparib for both ECM compositions which highlights the significance of synergy between genetic alterations and ECM landscape of lung cancer cells in regulation of therapeutic response. Furthermore, in alignment with the drug proximity analysis, treatment of cells with erlotinib showed an ECM-independent response and almost half of the population was eliminated with the given dose for both cell lines (Figure 6f). Together, these findings indicate an unprecedented potential of network-based drug efficacy screening for prediction of therapeutic response while considering the mutational background and tumor ECM composition of patients.

## DISCUSSION

While the ECM is among the key modulators of the TME, its precise contributions to cancer heterogeneity and therapy response are still not fully elucidated. In this study, we aimed to unveil how ECM influences oncogenic processes through latent ECM-mediated signaling pathways that extend beyond immediate matrix components, to demonstrate the prognostic and therapeutic potential of ECM-driven tumor stratification. Therefore, we integrated transcriptomic, proteomic, and phosphoproteomic profiles of matrisome and showed that LUAD tumors can be robustly classified into four distinct ECM barcode clusters, each associated with unique transcriptomic and network-level properties, and differential therapeutic vulnerabilities. We observed that these grades represent ECM characteristics, both through the ECM scores and independently determined highly variable genes. In our approach, we assessed change in ECM expression of a patient’s tumor compared to matching normal sample, which enabled us to account for patient-specific ECM heterogeneity.

We observed that higher ECM grades had elevated stromal content and reduced tumor purity. Thus, elevated ECM enrichment provides an environment not only suitable for the growth of tumor cells but also stromal cells such as CAFs^22^, which upon recruitment may take part in tumorigenesis and metastasis. CAFs are among the main sources of ECM deposition. Their reciprocal relationship with the surrounding ECM makes them critical targets for maintaining ECM homeostasis. In this study, we successfully showed increased CAF enrichment in higher ECM grades using cellular deconvolution methods from bulk transcriptomic data. Considering the inherent heterogeneity of CAFs, this finding could further be dissected with the use of higher-resolution technologies, such as single-cell RNAseq, to detect other subpopulations and cell states of CAFs that are influenced by altered ECM characteristics^72^.

We unveiled a gradual increase in EMT, invasiveness, CSC population and integrin activation along with higher risk of recurrence from ECM-G1 to ECM-G4. Significantly higher short-term recurrence probability in the ECM-G4 group can be attributed to the maintenance of CSCs as they drive differentiation and self-renewal^73,74^. These findings reveal that different ECM profiles are associated with distinct cellular processes and clinical features that align consistently with each other.

We also demonstrated the influence of ECM on KRAS-EGFR mutation exclusivity. We observed that a significant number of patients with lower ECM grades had EGFR mutations, and patients with higher ECM grades exhibited more KRAS mutations. EGFR is known to act in synergy with integrins and take part in cell-ECM interactions, probing for matrix stiffness^75,76^. Given the interplay between ECM and EGFR, higher ECM grades, likely showing increased ECM deposition, may already activate EGFR, resulting in more aggressive tumors. In contrast, tumor maintenance in lower ECM grades may rely mainly on EGFR mutations, with less support from ECM-mediated EGFR activation and thus resulting in milder tumors.

ECM grade-dependent alterations in pathways, clinical features, and mutation profiles prompted us to unravel the underlying causality of altered ECM profiles, which can be systematically explored through network modeling approaches. Previous studies have created matrisome co-expression networks^28,29^, yet to our knowledge, this study is the first to model the alteration in intracellular signaling in different ECM profiles in an integrative, patient-specific manner. Our matrisome-based focus on network modeling enabled us to go beyond the investigation of ECM signatures and reveal latent molecular players and hidden pathways taking part in ECM heterogeneity. We clearly showed that these networks could preserve patient heterogeneity but also were representative of the ECM components since the most enriched pathways were related to ECM interactions for all the constructed networks. We also revealed consistent patterns in pathways with both our previous findings and literature, such as EMT, KRAS signaling, angiogenesis, and the PI3K pathway, in line with its great role in mechanotransduction^77^.

Despite the explicit role of ECM in cancer cell behavior, the reciprocal interaction between ECM composition and genetic profile has not yet been fully uncovered and translated into the development of patient-specific therapeutic approaches. Currently, various ECM components have been shown as attractive therapeutic targets, particularly fibrillar collagens and hyaluronic acid (HA) as well as enzymes involved in ECM remodeling^36^. However, clinical trials have shown that ECM-directed therapies have highly context-dependent regulation of response and require comprehensive evaluation of genetic background of disease alongside with the characteristics of the specific TME^78^. Here, we constructed ECM grade-specific consensus networks, which exhibited distinct properties for ECM grades of patients while also preserving shared core proteins. Importantly, beyond selecting immediate ECM components as targets, we identified latent drug targets and pathways involved in ECM remodeling via network modeling. These findings support that the potency of targeted therapies are not only shaped by intercellular molecular signatures but also ECM-induced signaling.

We validated our hits using KRAS- and EGFR-mutated lung adenocarcinoma cell lines cultured on different ECM compositions, based on our observation that ECM alterations shape the mutational landscape of patients, specifically the KRAS-EGFR exclusivity. Our network-based drug screening identified specific associations of PARP1 and PARP2, targets of olaparib, with ECM-G1, while the EGFR inhibitor erlotinib has been found proximate to all ECM grades. Although olaparib is mainly used for treatment of BRCA1/2 mutated ovarian and breast cancers, benefits in metastatic NSCLC have been under investigation^79–81^. Notably, one recent preclinical study found that olaparib specifically induced apoptosis in BRCA1/2 depleted NSCLC cells due to deficiency in homologous recombination^82^. In alignment with our *in silico* findings, experimental results validated the stronger effect of olaparib on EGFR-mutated H1975 cells when they were seeded onto healthy lung-derived ECM compared to tumor-derived rBM. Intriguingly, this result was specific to EGFR mutated cells, and we did not observe a significant difference between two ECM compositions for KRAS-mutated A549 cells emphasizing the substantial impact of genetic drivers on matrix-guided regulation of drug response. In the second group, we confirmed that the response outcome to erlotinib (EGFR inhibitor) was independent from the ECM composition for both cell lines. Erlotinib targets constitutively active EGFR by competing with ATP binding that inhibits its autophosphorylation and hinder downstream proliferative signaling. Altered sensitivity to erlotinib could be driven by EGFR on-target mutations or activation mutations on adjacent signaling routes^83^. Together with experimental validation, we presented a computational model that successfully predicts the potency of drugs in proximity with ECM characteristics.

In this study, we performed early integration of the multi-omic data of the matrisome and stratified the patients accordingly. Our early integration approach provides a foundation upon which late integration strategies, including leveraging graph neural networks, can be constructed to delineate the complexity and topology of biological networks. Explainability of these models can also guide us to unravel other hidden patterns and potential therapeutic targets^84,85^.

A major limitation in patient stratification studies is the incomplete multi-omic data across patients, which results in restricting the analysis to only those samples containing all omic layers when applying integrative approaches. This applies to our study as well, compounded by the lack of tumor samples matched with normal tissue, leaving us with 101 LUAD samples. This issue can be tackled with larger patient cohorts with multi-omic data or approaches that utilize partial graph fusion mechanisms where each omic representation is iteratively updated with other omic data from overlapping patients while also keeping patients with partial data^86^.

Our approach to finding the intracellular networks behind altered ECM characteristics shared by the ECM grades revealed ECM-dependent drug candidates that may not directly target the ECM or its regulators but target intermediate proteins of the associated latent pathways. This suggests that ECM-based therapeutic strategies may benefit from considering also the ECM-induced intracellular signatures. Therefore, we expect that applying ECM-based stratification techniques to a pan-cancer cohort could be highly impactful for designing optimal treatment strategies, combining tumor molecular profiles and ECM-specific features.

## METHODS

### Data Acquisition

The dataset analyzed in this study was retrieved from the CPTAC-3 via the Proteomic Data Commons (PDC), comprising matched tumor and normal adjacent tissue samples from 101 patients with LUAD, along with multi-omic and clinical annotations. Transcriptomic data contains upper-quartile normalized and log₂-transformed RPKM values for each gene across all samples, and protein abundance and phosphorylation data contain log₂-transformed tandem mass tag (TMT) values^87^. Gene expression values were normalized by z-scores transformation across tumor samples prior to downstream analyses.

The Matrisome Project database was used to obtain the proteins of matrisome categories and components; the core matrisome (195 ECM glycoproteins, 44 collagens, 35 proteoglycans) and matrisome-associated genes (238 ECM regulators, 344 secreted factors, 171 ECM-affiliated) ^24,25^.

### Score Calculation for Gene Groups

In this study, we employed a unified scoring function based on the average expression of a given gene or protein set. This approach was applied across multiple contexts—including component-level scores, pathway activity scores, and network-based scores—where each context provides a biologically meaningful set of features. For example, a network score reflects the mean expression of all nodes within a molecular network, while a pathway score captures the average expression of genes involved in a specific biological pathway. This framework enables consistent and interpretable quantification across diverse molecular contexts.

Given a set of genes, 𝑁_𝑖_, the 𝑆𝑐𝑜𝑟𝑒(𝑁_𝑖_) is calculated via Equation 1.

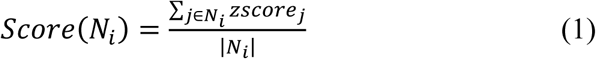

Cancer-associated gene sets for EMT, CSC, invasiveness, and integrin scores were retrieved from our previously published work and calculated using Equation 1^16,88^.

### ECM-Based Patient Barcoding and Consensus Clustering

Patients were barcoded as six-dimensional score vectors, with each dimension corresponding to ECM categories: collagens, proteoglycans, glycoproteins, regulators, secreted factors, and affiliated genes. For each ECM category, the score was calculated using Equation 1, replacing z-score with fold change values. Specifically, transcriptomic data was filtered to include only the matrisome genes, and the expression level of NAT samples was subtracted from the expression level of tumor samples for each patient, representing the upregulation or downregulation of an ECM gene in the patient’s tumor sample compared to the patient’s NAT sample. Then, genes having a difference value below - 2 or above 2 were retained to get highly dysregulated ECM genes which were then grouped by their ECM categories. The resulting gene sets were used for computing patient score vectors. Protein abundance and phosphorylation data were processed the same way. Finally, transcriptomic, proteomic, and phosphoproteomic ECM scores were combined by taking the average to get a multi-omic ECM score vector which indicates the increase or decrease in gene expression, protein abundance, and phosphorylated protein abundance of the ECM categories in a patient-specific manner. These barcodes are unique for each patient and provide a holistic view of the regulation profile of the ECM components.

Next, patient barcodes were merged to construct a 6 𝑥 𝑛 matrix, where the columns represent patients, and the rows represent ECM categories. Monte Carlo reference-based consensus clustering (M3C) package was used to cluster the patients by setting the objective function to the proportion of ambiguous clustering (PAC), the clustering algorithm to k-means clustering, and the rest of the arguments to default^89,90^.

### Highly Variable Gene Identification

To observe genes distinctive of the ECM grades and show that these grades indeed represent ECM characteristics of the groups, genes with high variability of expression were identified using tumor z-scores. Genes with z-scores above 1 (high expression) or below - 1 (low expression) were counted separately and those with occurrence in at least 13 patients, except for ECM-G4 where the threshold for occurrence was in at least 8 patients, in either high or low expression were determined as highly variable genes.

### Cellular Heterogeneity

For detecting cellular heterogeneity, the R package immundeconve^91^ was used with RPKM values transformed into TPM, selecting the tools MCP-Counter^92^ and EPIC^93^. Estimation of STromal and Immune cells in MAlignant Tumor tissues using Expression (ESTIMATE) stromal scores and tumor purity values of tumors were already provided in the CPTAC clinical data^94^.

### Clinical Data Analysis

Survival and recurrence-free analyses were performed on ECM grades using the survival R package, and the log-rank test was done to calculate p-values ^95^. Short-term survival and recurrence-free analyses were performed within the first 600 and 300 days, respectively. Patients with events that occurred after these time points were assumed to be survivors or recurrent-free until that point. In addition to the log-rank test, a multivariate COX proportional hazards model was employed to calculate hazard ratios (HR) by considering age, sex, smoking status, and ECM grade.

### Transcription Factor Enrichment Analysis

TFs targeting the highly dysregulated ECM genes were found using TRRUST, where the significant regulators are determined with Fisher’s exact test (q-value<0.1)^96^. For each ECM grade, a TF expression score was calculated by taking the median expression of the TF from tumor z-score data. Among the significant TFs, the ones enriched in more than 25% of the corresponding ECM grade tumors in at least one ECM grade were retained. The set of TFs having a monotonically increasing or decreasing trend across ECM grades were used for visualization.

### Network Reconstruction

ECM-guided integrative network modeling for each patient was performed with the forest module of OmicsIntegrator2 which solves prize-collecting Steiner Forest (PCSF) problem on a given set of terminal proteins/genes and a reference confidence weighted PPI interactome. The final optimal PCSF contains terminal nodes connected either directly or via intermediate nodes by high confidence edges.

Given 𝐺(𝑉, 𝐸, 𝑐(𝑒), 𝑝′ (𝑣)) is a directed or undirected graph where 𝑉 is the node set, 𝐸 is the edge set, 𝑐(𝑒) > 0 is the cost for each edge 𝑒 ∈ 𝐸, and 𝑝′(𝑣) is the attenuated prize value for each node 𝑣 ∈ 𝑉.

The PCSF algorithm aims to find a forest 𝐹(𝑉_𝐹_, 𝐸_𝐹_) by minimizing the objective function in Equation 2.

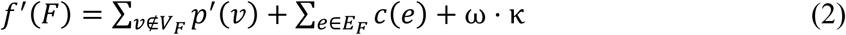

The first element calculates the sum of prizes of terminals that were not included in the resulting forest.

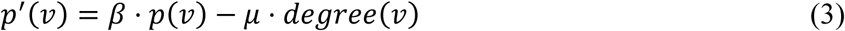

Where, 𝑝(𝑣) is the prize for node 𝑣, and 𝑑𝑒𝑔𝑟𝑒𝑒(𝑣) is the number of first neighbors of node 𝑣. 𝛽 is a parameter for tuning for the number of terminals to be included in the solution, and µ is a parameter adjusting to penalize the nodes based on their degrees.

The second element is the sum of edge costs in the resulting forest. The edge costs 𝑐(𝑒) is calculated from the confidence values 𝑐𝑜𝑛𝑓(𝑒) with Equation 4.

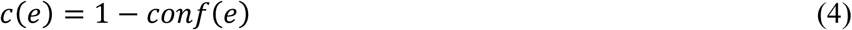

The third element is involved in enabling multiple trees to form in the solution by introducing an artificial node connected to terminals with an edge cost 𝜔. 𝛽, 𝜔, and 𝐷 parameters are tuned to reach optimal solutions. The 𝐷 parameter tunes for the maximum path length between terminals and the artificial node^97,98^.

Patient-specific signaling networks were constructed with terminal lists that included dysregulated ECM phosphoproteins and key regulators. The weighted PPI reference interactome used in network modeling was iRefIndex version v14^99^ (182,002 interactions between 15,759 proteins) where MI scores were the weights and the proteins “UBC”, “APP”, “ELAVL1”, “SUMO2“, and “CUL3” were removed^100^. The TF prizes were calculated by taking the average of the absolute fold change values of the highly dysregulated target ECM genes. For the highly dysregulated phosphoproteins, the highest absolute difference values were assigned as the prizes. For 𝛽, 𝜔, and 𝐷, values [2, 3, 4, 5] were used in combination, and 64 networks in total were constructed. Lastly, the final patient-specific networks were constructed by merging the two top ranking networks that had the minimal number of components and nodes and maximal terminal coverage.

### ECM Grade Specific Consensus Network Construction

For each grade, a consensus network was constructed from patient-specific networks. frequency of each edge within the patient population of an ECM grade was determined, and those observed in 30% of the patients were retained. Final consensus networks were obtained by selecting the largest connected component from the edge-cleared data. To have comparable node coverage across grades, edges were pruned such that grade-specific node counts remained approximately balanced. To account for variability in patient group sizes, only edges observed in at least four patients were retained. Consensus networks were then compared across grades using the Jaccard index.

### ECM Grade Specific Pathway Enrichment Analysis

Grade-specific pathway analysis was performed using the EnrichR^101^. Gene sets used in functional enrichment analysis were pathways from the Kyoto Encyclopedia of Genes and Genomes (KEGG) and REACTOME, hallmarks from the Molecular Signatures Database (MSigDB) and biological processes from Gene Ontology^102,103^. Functional enrichment analysis was performed using the nodes of patient-specific networks and pathways with an adjusted p-value below or equal to 0.05 and overlapping gene counts above 2 were considered enriched. These were then subjected to grade-wise trimming mentioned in the TF enrichment analysis part. Then, a pathway score was calculated for each patient using tumor z-score data by taking the average of the z-scores of genes related to the pathways. A similar score was calculated for each grade by averaging the pathway scores of patients of the same grade.

### Network Proximity Analysis for Drug Ranking

To identify candidate drugs associated with the ECM grades, we performed network proximity analysis. We computed the average minimal distances between the targets of each drug and the proteins in the ECM grade specific consensus networks. The same reference interactome used in network modeling was employed for drug proximity analysis.

Given a set of grade-specific proteins 𝑆 and drug target proteins 𝑇, the shortest path length 𝑑(𝑠, 𝑡) is calculated for each pair of drug targets, 𝑡, and grade-specific proteins, 𝑠, and the minimum length is picked for the pair and averaged on the drug target proteins to give the closest distance 𝑑_𝑐_(𝑆, 𝑇) as shown in Equation 5.

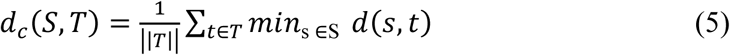

Then a reference distribution of closest distances is calculated by selecting two sets with the same size and degree as the original disease and drug target proteins 1000 times, and then the mean 𝜇_𝑑(𝑆,𝑇)_ and standard deviation 𝜎_𝑑(𝑆,𝑇)_ of the randomly generated distance values are used to calculate a relative proximity 𝑧(𝑆, 𝑇) as shown in Equation 6.

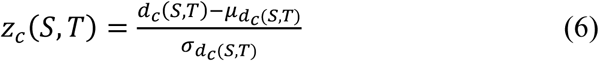

Drugs were considered proximal to the disease proteins when 𝑧_𝑐_ ≤ −0.15 and significantly proximal when 𝑧_𝑐_ ≤ −2.

Network proximity analysis was performed between the proteins of each ECM grade-specific consensus network and the targets of approved drugs (retrieved from the DrugBank^104^ database) using the source code by Guney et al. with default settings^33^. Drugs with corresponding proximity values were subset into the list of drugs in Oncotarget^66^.

### Statistical Analyses

Spearman rank correlation was performed for all correlation analyses. A two-sided Mann-Whitney U test was employed to calculate the p-values when comparing the distributions of ECM grades for the calculation of gene group scores, Holm–Bonferroni method was used for adjustment for multiple comparisons. Enrichment analyses were performed using Fisher’s Exact test. P-values smaller than or equal to 0.05 were considered significant throughout the study.

### Cell Culture and Drug Response

Human lung adenocarcinoma cell lines A549 (RRID: CVCL_0023) and H1975 (RRID: CVCL_1511) were obtained from American Tissue Culture Collection (ATCC) in March 2021 and cultured in growth medium DMEM/F12 (Lonza) and RPMI (Capricorn), respectively, supplemented with 10% fetal bovine serum (FBS) (Biowest) and 1% penicillin-streptomycin (P/S) (Gibco). All cultures were routinely tested and confirmed to be contamination-free. Drug response experiments were performed on a 96-well plate. Cultrex (R&D Systems) and decellularized lung (dLung) matrices were used as ECM cues. dLung matrix was produced according to our published protocol^70^. Briefly, healthy bovine lung tissue was minced into small pieces and cellular content was removed with extensive washes and freeze/thaw cycles. Decellularized tissue pieces were freeze-dried, cryo-milled, and thermal gelation was performed after pepsin digestion and neutralization. The surface of the wells was coated with either Cultrex or dLung hydrogel precursor solutions with 0.5 mg/ml ECM concentration. Precursor solutions were allowed to thermally crosslink at 37 °C and 5% CO2 within a cell culture incubator for 90 minutes. After crosslinking, cells were seeded onto the wells at a density of 3000 cells/well. Cells were allowed to attach onto ECM-coated surfaces for 16 hours. Then, the growth medium was aspirated, and cells were treated with erlotinib (2 µM, 10 µM and 50 µM) or olaparib (10 µM, 50 µM, 100 µM). A control group with 1:100 DMSO concentration was included to calculate cell viability at the end point. Cells were treated with the drugs for 3 days at given concentrations and their metabolic activities were assessed with CellTiter Glo (Promega) assay as a measure of cell viability. Cellular viability was calculated in percentage relative to the control group to observe the efficacy of drugs in Cultrex and dLung-coated surfaces. Three independent replicates were used, and statistical significance was tested with unpaired t-test.

## Code Availability

All the code used for data analysis and generation are available at: https://github.com/adnsk22/ecm_grades. ECM-based consensus clustering was performed using Monte Carlo Reference-based Consensus Clustering accessible at https://www.bioconductor.org/packages/release/bioc/html/M3C.html. ECM-guided patient-specific network modeling was performed using Omics Integrator 2 which is available at https://github.com/fraenkel-lab/OmicsIntegrator2. All functional enrichment analyses were performed using enrichR available at https://cran.r-project.org/package=enrichR. All cellular deconvolution analyses were performed using immundeconv accessible at https://github.com/omnideconv/immunedeconv. Network-based drug screening was performed using the script available at https://github.com/emreg00/toolbox.

## Data Availability

The CPTAC LUAD dataset is available from the CPTAC Portal: https://pdc.cancer.gov/pdc/study/PDC000153. The list of matrisome proteins are available in The Matrisome Project: https://sites.google.com/uic.edu/matrisome/matrisome-annotations/homo-sapiens?authuser=0. EMT, INV and CSC gene lists were retrieved from the supplementary material of Kuşoğlu et al.: https://doi.org/10.1002/advs.202309966. Transcription factors were retrieved from TRRUST v2: https://www.grnpedia.org/trrust/Network_search_form.php. The reference PPI network, iRefIndex v14 was downloaded from Omics Integrator git repository: https://github.com/fraenkel-lab/OmicsIntegrator2. Drug molecules and their target proteins were downloaded from DRUGBANK: https://go.drugbank.com/. Oncotarget drug list was retrieved from the supplementary material of Mundi et al.: https://doi.org/10.1158/2159-8290.CD-22-1020

## Acknowledgements

NT was supported by the Research Projects Funding Program of TUBITAK under the project number 121E245 and the National Leader Researchers Program of TUBITAK under the project number 121C292.

## Competing interests

The authors declare no potential conflicts of interest.

## Authors’ contributions

Conceptualization: EÖ, NT

Data curation: AD, NT

Formal analysis: AD, NT

Methodology: AD, EÖ, NT

Conducting experimental validation: SS, EÖ

Supervision: EÖ, NT

Visualization: AD, SS

Writing – original draft: AD, SS, EÖ, NT

Writing – review and editing: AD, SS, EÖ, NT

